# *Bifidobacterium breve* UCC2003 induces a distinct global transcriptomic programme in neonatal murine intestinal epithelial cells

**DOI:** 10.1101/2020.03.27.011692

**Authors:** Raymond Kiu, Agatha Treveil, Lukas C. Harnisch, Shabhonam Caim, Charlotte Leclaire, Douwe van Sinderen, Tamas Korcsmaros, Lindsay J. Hall

## Abstract

*Bifidobacterium* is an important gut microbiota member during early life that is associated with improved gut health. However, the underlying health-driving mechanisms are not well understood, particularly how *Bifidobacterium* may modulate the intestinal barrier via programming of intestinal epithelial cells (IECs). In this study, we sought to investigate the global impact of model strain *Bifidobacterium breve* UCC2003 on the neonatal IEC transcriptome, including gene regulation and pathway modulation. Small IECs from two-week-old neonatal mice administered *B. breve* UCC2003 for three consecutive days or PBS (control group) were subjected to global RNASeq, with various bioinformatic approaches used to determine differentially expressed genes, pathways and affected cell types between control and experimental groups. Whilst colonisation with *B. breve* had minimal impacts on the neonatal microbiota, we observed extensive regulation of the IEC transcriptome; ~4,000 genes significantly up-regulated, including key genes associated with epithelial barrier function. Enrichment of cell differentiation and cell proliferation pathways were observed, along with an overrepresentation of stem cell marker genes, indicating an increase in the regenerative potential of the epithelial layer. Expression of distinct immune-associated pathway members (e.g. Toll-like Receptors) were also affected after neonatal *B. breve* colonisation. In conclusion, *B. breve* UCC2003 plays a central role in driving universal transcriptomic changes in neonatal IECs that enhances cell replication, differentiation and growth, predominantly in the stem cell compartment. This study enhances our overall understanding of the benefits of *B. breve* in driving intestinal epithelium homeostatic development during early life, with potential avenues to develop novel live biotherapeutic products.

## Introduction

*Bifidobacterium* represents a keystone member of the early life gut microbiota [1-3]. Certain species and strains are found at high levels in vaginally delivered breast-fed infants including; *Bifidobacterium longum* subsp. *infantis, B. longum* subsp. *longum, B. bifidum, B. pseudocatenulatum* and *B. breve* [4-7]. As a dominant member of the neonatal gut microbiota, *Bifidobacterium* is associated with metabolism of breast milk, modulation of host immune responses, and protection against infectious diseases [8-11]. However, the mechanisms driving improved health outcomes during early life are largely underexplored.

A key interface between *Bifidobacterium* and the host is the intestinal epithelial cell (IEC) barrier [12, 13]. Previous studies have indicated that certain strains of *Bifidobacterium* specifically modulate IEC responses during inflammatory insults, which may help protect from certain gut disorders [14-16]. In murine experimental models, previous work by our group has shown that infant-associated *B. breve* UCC2003 modulates cell death-related signalling molecules, which in turn reduces the number of apoptotic IECs [17]. This protection from pathological IEC shedding appeared to be via the *B. breve* exopolysaccharide (EPS) capsule and the host-immune adaptor protein MyD88. Another strain of *B. breve,* NumRes 204 (commercial strain) has also been shown to up-regulate the tight junction proteins Claudin 4 and Occludin in a mouse colitis model [18, 19].

Many of the studies to date have focused on the role of *Bifidobacterium* and modulation of IECs in the context of acute or chronic gut inflammation, with expression profiling limited to specific immune or apoptosis signalling targets [14, 20-22]. As many of these studies have involved pre-colonisation of the gut with *Bifidobacterium* strains, followed by inflammatory insult, this suggests that initial priming during normal ‘healthy’ conditions may modulate subsequent protective responses. Furthermore, these studies have often been performed in adult mice rather than exploring effects during the early life developmental window, where *Bifidobacterium* effects are expected to be most pronounced. Previous work has indicated that there is significant modulation of the neonatal IEC transcriptome in response to gut microbiota colonisation, but to date no studies have probed how particular early life associated microbiota members, like *Bifidobacterium* may modulate neonatal IEC responses [23]. Thus, to understand if and how *Bifidobacterium* may modulate IEC homeostasis during the early life developmental window, we colonised neonatal mice with *B. breve* UCC2003 and profiled transcriptional responses in isolated small intestine IECs using global RNA-Seq. Our analysis indicated whole-scale changes in the transcriptional programme of IECs (~4,000 significantly up-regulated genes) that appear to be linked to cell differentiation/proliferation and immune development. Notably the stem cell compartment of IECs seemed to elicit the strongest gene signature. These data highlight the role of the early life microbiota member *B. breve* UCC2003 in driving early life epithelial cell differentiation and maturation; impacting intestinal integrity and immune functions, which provides a mechanistic basis for understanding associated health-promoting effects.

## Results

To examine the effects of *B. breve* UCC2003 on the transcriptional profiles of host IECs under homeostatic conditions, we extracted RNA from isolated IECs of healthy two-week old neonatal mice (control group) and mice gavaged with *B. breve* UCC2003 for three consecutive days (*n*=5 per group). Isolated RNAs from IECs were subjected to RNA-Seq to determine global mRNA expression (Fig. 1). Subsequently, Differential Gene Expression (DGE) analysis was performed to understand *B. breve*-associated gene regulation

**Fig. 1.**
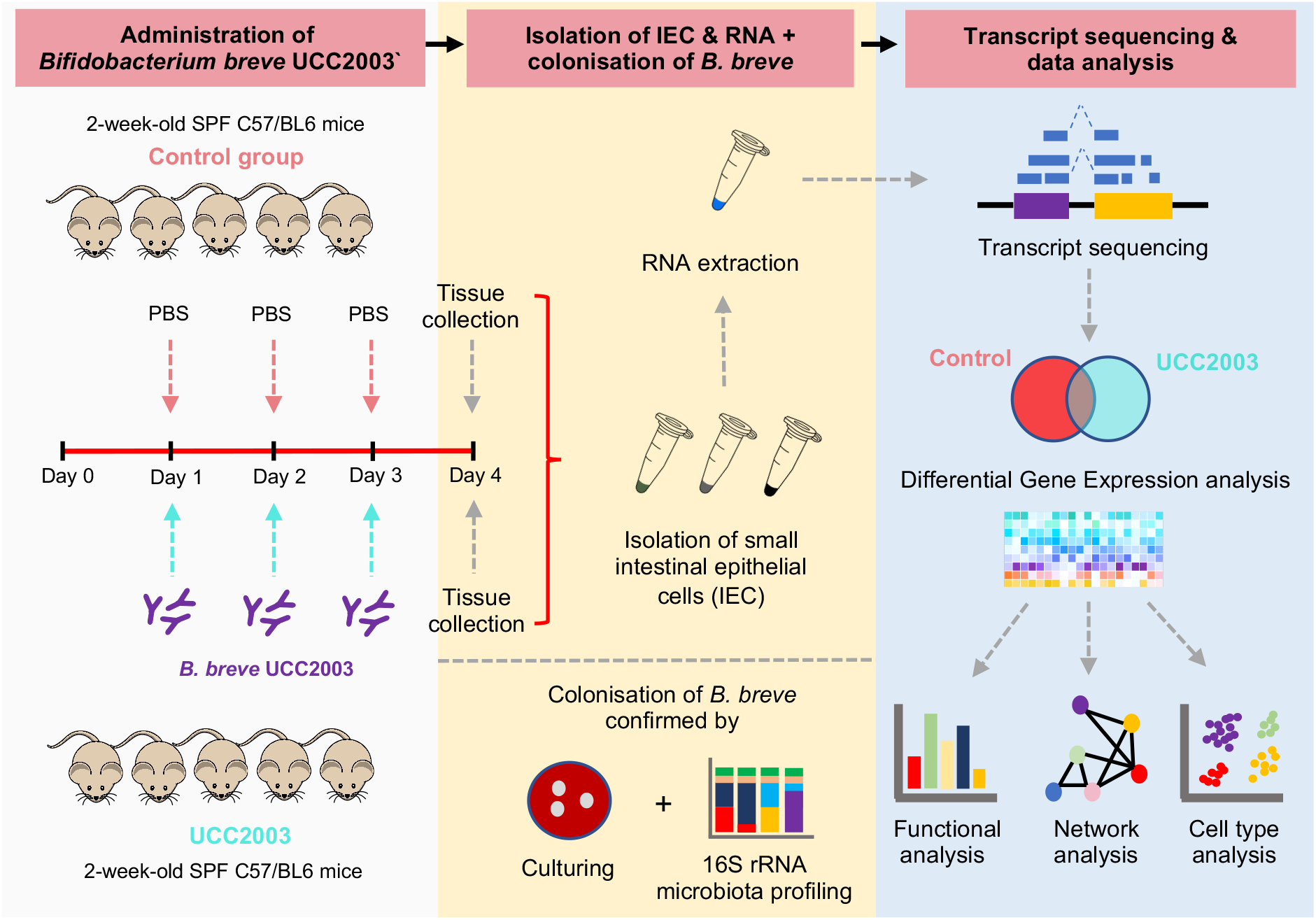
Schematic representation of study design, experimental validation and *in silico* analysis workflow.

### Colonisation of *B. breve* UCC2003 and impact on the wider neonatal microbiota

Initially, we confirmed gut colonisation of *B. breve* UCC2003 and impact on the wider microbiota using culture and 16S rRNA microbiota profiling approaches (Fig. 2a-b). We observed high levels of *B. breve* UCC2003 across the four days in faecal samples, with higher levels of *B. breve* UCC2003 within the colon (~10^8^ CFU/g), when compared to the small intestine (~10^5^ CFU/g; Fig. 2b). Based on 16S rRNA analysis, relative abundance of *Bifidobacterium* increased significantly in the UCC2003 group (*P*=0.012) following bacterial administration, while the control group displayed very low relative *Bifidobacterium* abundance (~0.01%; Fig. 2c). Principal component analysis (PCA) on gut microbiota profiles (control vs UCC2003) showed a distinct change in microbial community composition in the UCC2003 group; primarily driven by *Bifidobacterium*, and to a lesser extent by *Lactobacillus* and *Bacteroides* (Fig. 2d). Supplementation of *B. breve* also significantly increased the overall microbial diversity in the UCC2003 group (Fig. 2e). Further Linear Discriminant Analysis (LDA) demonstrated that although *Bifidobacterium* was enriched in UCC2003 group, colonisation had minimal impact on overall microbiota profiles, although very low relative abundance (<2%) microbiota members *Streptococcus, Ruminococcus, Prevotella, Coprococcus* were significantly reduced in the *B. breve* UCC2003 group (Fig. 2f-g).

**Fig. 2.**
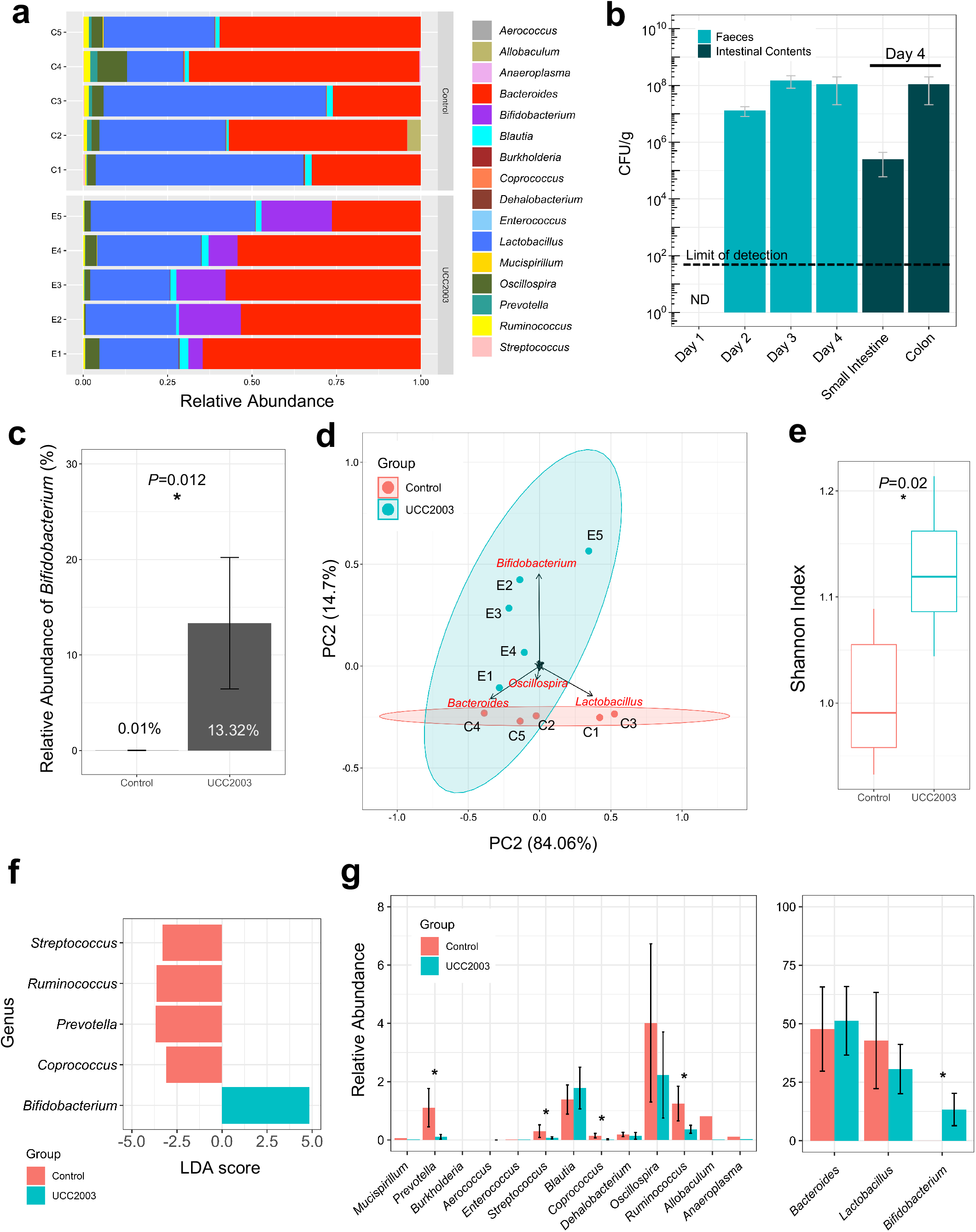
16S rRNA amplicon sequencing analysis of murine intestinal microbiota. (a) 16S rRNA gene profiling of mice gut microbiota at genus level on Day 4. (b) Dynamics of *B. breve* UCC2003 load (CFU/g) from Day 1 (prior to *B. breve* administration) through Day 4. *B. breve* was present in intestines throughout (small intestines and colon; on Day 4). (c) Relative abundance of genus *Bifidobacterium* in UCC2003 group is significantly increased. (d) Principal Component Analysis on mice gut microbiota. (e) Shannon diversity index on mice gut microbiota. Data are represented as mean ± SD. Significance test: *t*-test (\**P*<0.05; two-sided). (f) Linear Discriminant Analysis (LDA) showing enriched taxa in each group. (g) Relative abundance comparison in all genus. * *P*<0.05 in LDA.

### Impact of *B. breve* UCC2003 on the neonatal intestinal epithelial transcriptome

To understand the distribution of samples based on IEC gene expression profiles we performed PCA analysis (Fig. 3a; Table S1). All samples clustered according to group (control vs UCC2003), suggesting a significant impact of *B. breve* UCC2003 on gene expression profiles, with distance-wise clustering (Jensen-Shannon) also supporting separation of experimental groups (Fig. 3b). To define Differentially Expressed Genes (DEG), we employed a filter of absolute log2(fold change) > 1.0 (with adjusted *P* < 0.05), which equates to a minimum two-fold change in gene expression (Fig. 3c-e; Table S2). After analysis, a total of 3,996 DEGs were significantly up-regulated, while 465 genes were significantly down-regulated in *B. breve* UCC2003 supplemented animals when compared to controls (Fig. 3c and 4a). Notably, we also performed the same experimental protocol on healthy mice aged 10-12 weeks, and we did not observe any DEGs, suggesting *B. breve* UCC2003 modulation of IECs is strongest within the early life window under homeostatic conditions.

**Fig. 3.**
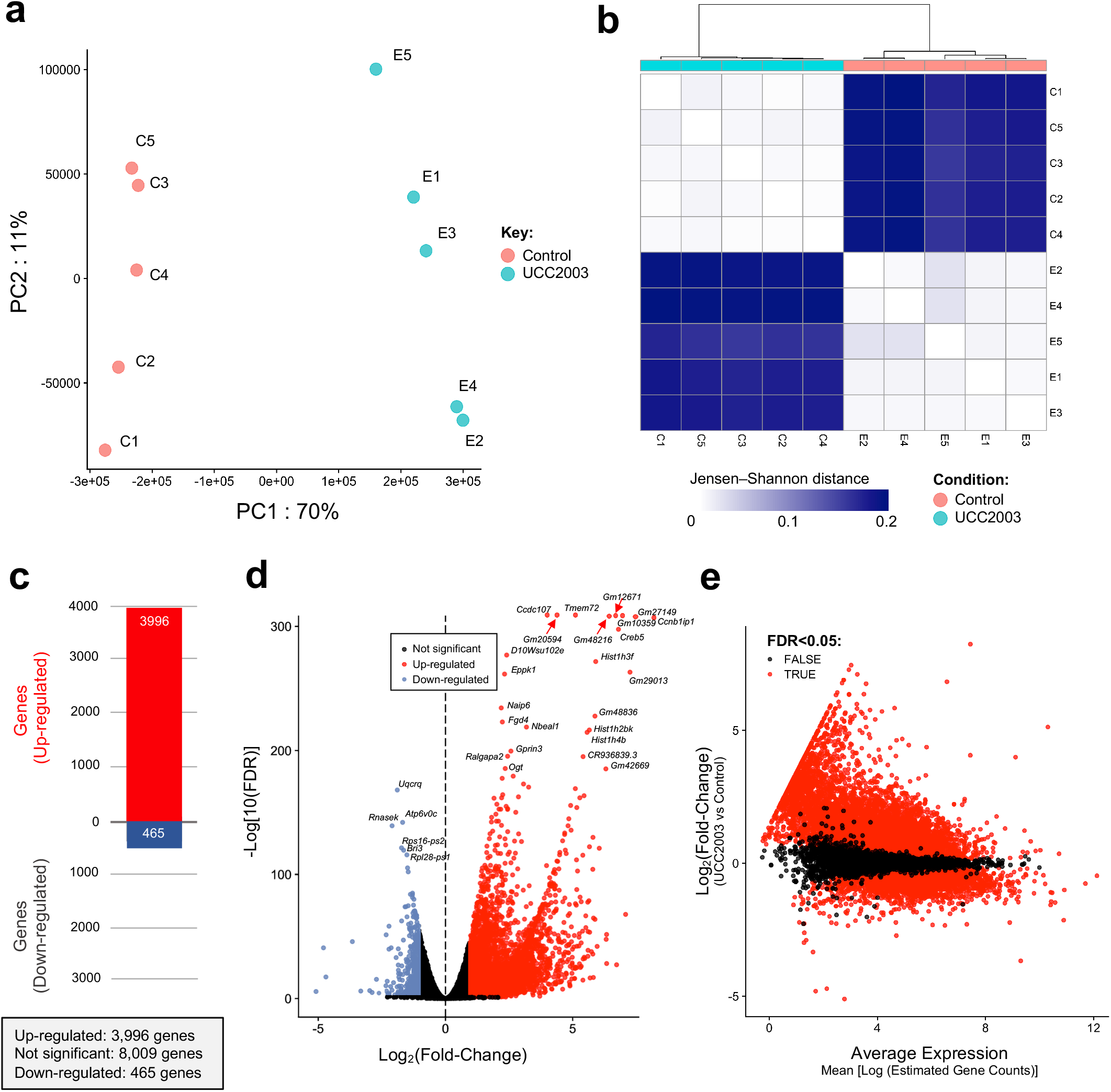
RNA-Seq analysis and statistics (a) Principal component analysis showing distinct overall gene expression profiles across all individual samples based on 12,965 highly-expressed genes; (b) Clustering of individual RNA-seq samples based on Jensen-Shannon distance; (c) Total number of differentially expressed genes (DEG) when comparing two conditions (UCC2003 vs Control), DEG with Log_2_FC>1.0 (up-regulation) or Log_2_FC <-1.0 (down-regulation) are considered as significantly regulated genes; (d) Volcano plot and (e) MA plot on global gene expression (UCC2003 vs Control). Genes that passed the significance filter (FDR <0.05) are labelled as red dots.

To determine the functional role of the DEGs, we examined the most significantly regulated genes ranked by False Discovery Rate (FDR) adjusted *p* values (or, *q* values). We first looked at the top 20 up-regulated DEGs in the *B. breve* UCC2003 experimental group (Fig. 4b). Most genes annotated with known biological processes were cell differentiation and cell component organisation functions including *Ccnb1ip1, Hist1h4b, Vps13b* and *Fgd4* (annotated in the PANTHER Gene Ontology [GO] Slim resource). Two genes were involved in cell death and immune system processes, namely *Naip6* and *Gm20594* (Table S3). When we ranked the top-regulated genes using log2-fold change, we observed increased expression of *Creb5*, which is involved in the regulation of neuropeptide transcription (cAMP response element binding protein; CREB) (Fig. 4c). CREB is also known to regulate circadian rhythm, and we also identified additional circadian-clock-related genes that were significantly up-regulated including *Per2* and *Per3*. We noted that several top down-regulated DEGs were annotated as genes involved in metal binding, or metal-related genes including *Mt1, Mt2, Hba-a1, Hbb-bt* and *Ftl1-ps1* (Fig. 4d; Table S4). These data suggest indicates *B. breve* stimulates specific genes involved in important physiological processes highlighting the importance of this microbiota member during the early life developmental window.

**Fig. 4.**
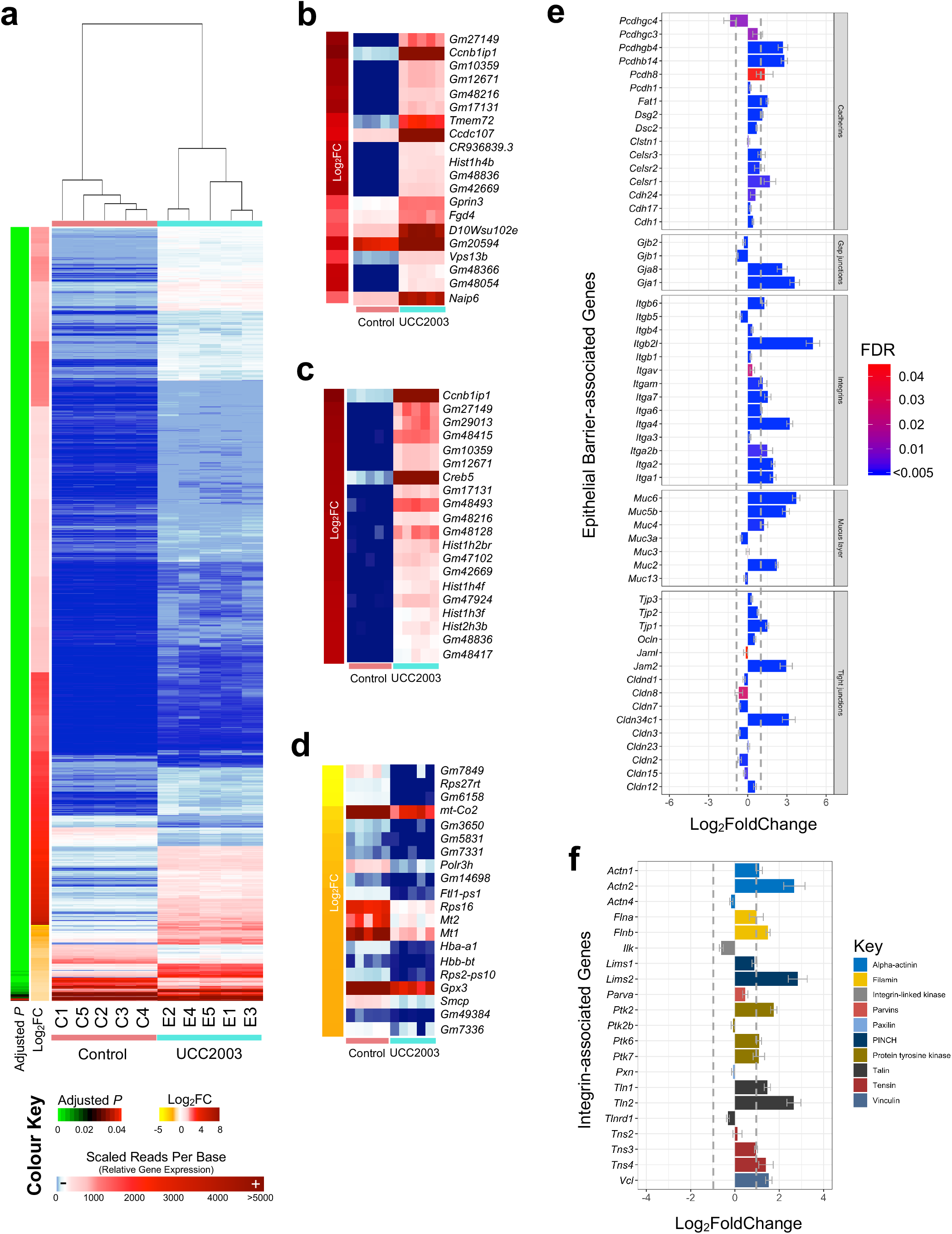
Gene expression analysis (a) Clustered normalised gene expression profiles on 4,461 significantly regulated genes (up- and down-regulated; FDR<0.05) in mice induced by *B. breve* UCC2003; (b) Top 20 significantly regulated genes ranked by FDR (q-value); (c) Top 20 significantly up-regulated genes ranked by Log_2_FC values; (d) Top 20 significantly down-regulated genes ranked by Log_2_FC values. Gene expression of (e) Epithelial integrity-associated DEG (FDR<0.05), and (f) Integrin-associated DEG in UCC2003 group were shown in the bar charts with dotted line indicating the threshold of significance (absolute Log_2_FC>1.0). Data are represented as Mean ± SE.

### *B. breve* UCC2003 modulation of intestinal epithelial barrier-associated genes

As *B. breve* strains have been previously shown to modulate certain tight junction/barrier-related proteins, we next investigated DEGs associated with intestinal epithelial barrier development/intestinal structural organisation (Fig. 4e). Several tight-junction (TJ) structural-associated DEGs were observed, including Claudin-encoding gene *Cldn34c1* (Log2 fold-change [LFC] 3.14), Junction Adhesion Molecules-encoding genes *Jam2* (LFC 2.9), and Tight Junction protein (also called Zonula Occludens protein; ZO) -encoding gene *Tjp1* (LFC 1.49). Other important TJ-associated protein-encoding genes including *Ocln* (encodes Occludin), and *Tjp2* and *Tjp3* and *Cldn12* (which represent ZO encoding genes) appeared to be transcriptionally up-regulated, although they did not pass the significant foldchange threshold. Genes that encode integrins (involved in regulation of intracellular cytoskeleton) also exhibited a trend of increased expression (13/14; 92.8%). Both Piezo genes, which assist in tight junction organisation, *Piezo1* (LFC 1.25) and *Piezo2* (LFC 1.9), were significantly up-regulated in the *B. breve* UCC2003 treated group.

Over 90% of cadherins, proteins associated with the assembly of adherens junctions (Fig. 4e) were up-regulated; including *Pcdhb14* (LFC 2.8), *Pcdhgb4* (LFC 2.7), *Pcdh8* (LFC 1.3), *Fat1* (LFC 1.5) and *Dsg2* (LFC 1.1). Interestingly, several genes (4/7; 57.1%) involved in mucus layer generation were significantly up-regulated in the UCC2003 experimental group including *Muc2* (LFC 2.2), *Muc6* (LFC 3.7), *Muc5b* (LFC 2.9), and *Muc4* (LFC 1.24). Genes *Gja1* (LFC 3.59) and *Gjb8* (LFC 2.63) that encode gap junction proteins were also up-regulated. In addition, we also investigated differential expression of genes associated with integrin assembly and downstream integrin signalling pathways (Fig. 4f). Over 70% (16/21) of these genes were up-regulated, with 52.3% (11/21) significantly increased in gene expression in the UCC2003 group (LFC >1.0).

We observed increased expression of genes associated with IEC barrier development including cadherins, gap junctions, integrins, mucus layer-associated genes, and several key tight junction proteins. These strongly induced gene expression profiles suggest that *B. breve* UCC2003 is involved in enhancing epithelial barrier development in neonates.

### *B. breve* UCC2003 modulates cell maturation processes

We next sought to understand the biological functions of up-regulated DEGs by employing PANTHER GO-Slim functional assignment, and process/pathway enrichment analysis (Fig. S1; Table S5-S9). DEGs were predominantly involved in general biological processes including cellular process (901 genes) and metabolic process (597 genes; Table S5). At the molecular function level, DEGs were primarily assigned to binding (868 genes) and catalytic activity (671 genes; Table S6), with Olfactory Signalling Pathway and Cell Cycle (biological) pathways also found to be enriched (Table S7).

To delve further into the data, we constructed a signaling network based on up-regulated DEGs (n=3,996) with the aim of identifying specific gene networks involved in important signalling pathways (Fig. 5a). Overall, 1,491 DEGs were successfully mapped (37.3%) to a signalling network that comprised 8,180 genes. Four individual clusters of genes were detected, with functional assignment and pathway analysis implemented on these clusters (Fig. 5a). All gene clusters were associated with cell differentiation and maturation, with cluster 1 (68 genes) linked specifically with DNA replication and transcription, cluster 2 (26 genes) with cell growth and immunity, cluster 3 (11 genes) with cell replication, and cluster 4 (72 genes) related to cell cycle and cell division (Table S10).

**Fig. 5.**
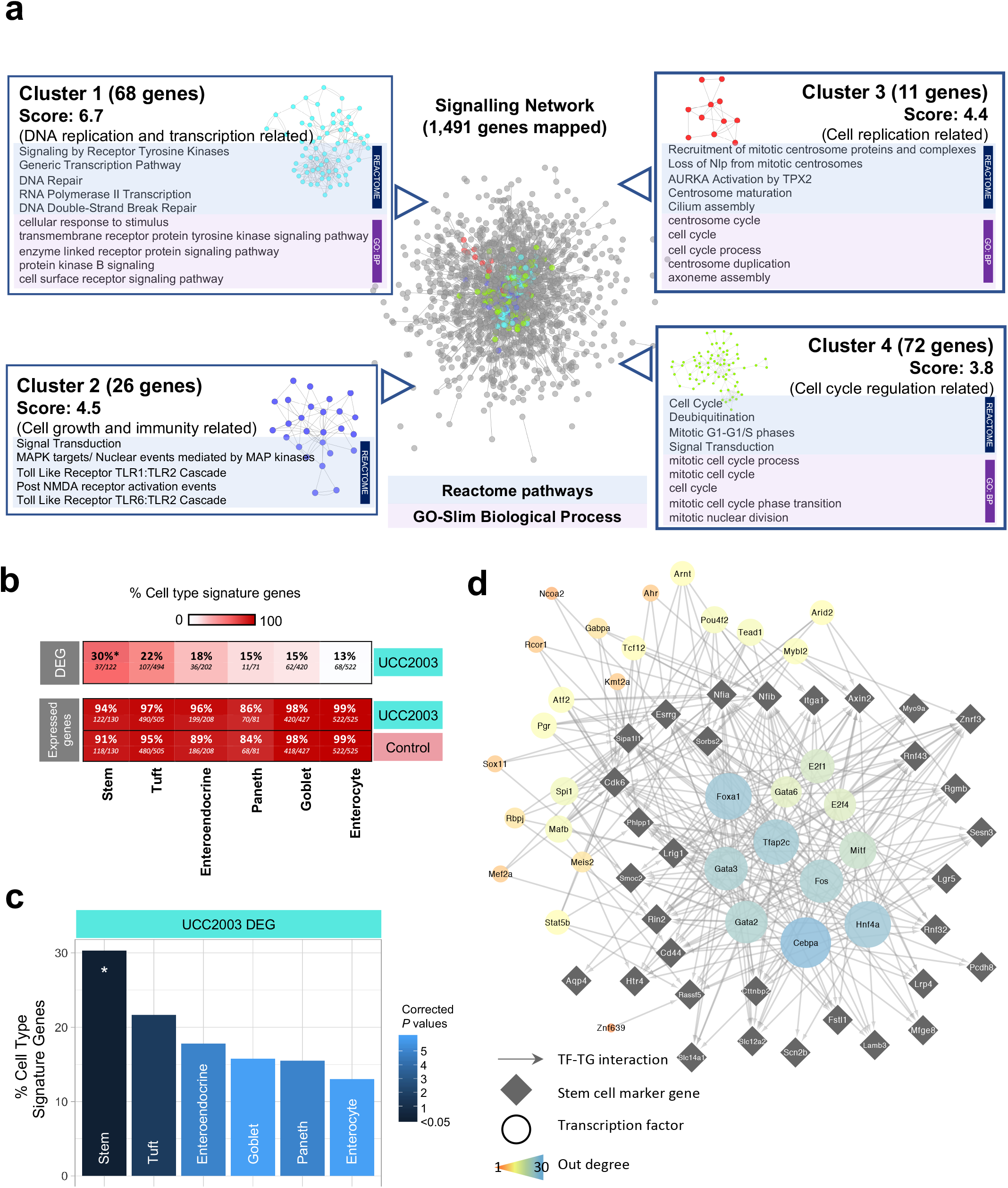
Signalling network analysis, IEC subtyping and key regulator analysis (a) Cluster analysis of signalling network for significantly up-regulated genes (*n*=3,996). Representative enriched pathways (Reactome) and GO terms (Biological Process) identified in each individual cluster were listed alongside. (b) Heat plot showing percentage of cell type signature genes in DEG and expressed genes (both control and UCC2003 groups). All expressed genes are well represented in IEC cell type signature genes. (c) Cell type analysis on IEC DEGs using known cell-specific signature genes. Stem cells were statistically overrepresented in DEGs. * *P*<0.05. (d) Key regulators of stem cell DEGs.

### Intestinal cell type analysis on DEGs identifies significant enrichment of epithelial stem cells

IECs include several absorptive and secretory cell types, namely enterocytes, Paneth cells, goblet cells, enteroendocrine cells, tuft cells and stem cells. As these cells perform different functions in the gut, it was important to understand whether *B. breve* UCC2003 had a cell type specific effect on the intestinal epithelium. Using known cell type specific gene markers [24], we identified cell type gene signatures modulated within the UCC2003 group (Fig. 5b-c). Importantly, all cell type markers were well represented in the expressed genes of the whole IEC transcriptomics data from both groups (control + UCC2003), thus validating the presence of all IEC types in our study data (Fig. 5b). Cell type analysis of genes differentially expressed after *B. breve* UCC2003 supplementation, revealed that stem cell marker genes were significantly enriched (30%; *P* < 0.05) among the six IEC types (Table S11). Signatures of other cell types were also present (linking to marker genes in the DEG list) but not significantly over-represented: Tuft cells (22%), enteroendocrine cells (18%), goblet cells (15%), Paneth cells (15%) and enterocytes (13%; Fig. 5c). These data indicated that intestinal epithelial stem cells, cells primarily involved in cell differentiation, were the primary cell type whose numbers and transcriptomic programme were regulated by *B. breve* UCC2003.

Further investigation of this stem cell signature revealed that of the 37 differentially expressed marker genes, 35 are up-regulated in the presence of *B. breve* UCC2003. This indicates an increase in the quantity of stem cells or semi-differentiated cells in the epithelium, consistent with the overrepresentation of cell cycle and DNA replication associated genes observed in the whole differential expression dataset. Functional analysis of the 37 stem cell signature genes revealed only one overrepresented process - Regulation of Frizzled by ubiquitination (*P* < 0.05), which is a subprocess of WNT signalling. WNT signalling is important in maintaining the undifferentiated state of stem cells [25].

Finally, we employed a network approach to predict key transcription factor (TF) regulators of the differentially expressed stem cell marker genes, through which *B. breve* UCC2003 may be acting (Fig. 5d). Using the TF-target gene database, DoRothEA, we identified expressed TFs known to regulate these genes [26, 27]. Five genes had no known and expressed regulator, thus were excluded. Hypergeometric significance testing was used to identify which of these TFs are the most influential (see Methods for details). This analysis identified 32 TF regulators (Fig. 5d). Of these regulators, 12 were differentially expressed in the IEC dataset (all up-regulated): *Fos, Gabpa, Rcor1, Arid2, Tead1, Mybl2, Mef2a, Ahr, Pgr, Kmt2a, Ncoa2* and *Tcf12*. Functional analysis of all the TF regulators and their targeted genes together, revealed overrepresented functions relating to WNT signalling, histone methylation for self-renewal and proliferation of hematopoietic stem cells and nuclear receptor (incl. estrogen) signalling (Table S12). These data provide evidence that *B. breve* UCC2003 directly affects key transcriptomic programmes regulating drives specific signalling processes, particularly within stem cells.

## Discussion

The early life developmental window represents a crucial time for microbe-host interactions that impacts health both in the short- and longer-term. Understanding how specific microbiota members modulate host responses during these life stages is crucial if we are to develop next-stage targeted microbiota therapies. Here we investigated how *B. breve* UCC2003 induces genome-wide transcriptomic changes in small intestine IECs of neonatal mice. We observed that *B. breve* had a global impact on the IEC transcriptome, evidenced by the large number of significantly up-regulated genes and pathways related to cell differentiation and cell proliferation, including genes associated with epithelial barrier function. We propose that *B. breve* is a key early life microbiota member driving fundamental cellular responses in IECs, particularly within the stem cell compartment, and thus drives epithelial barrier development and maintenance during neonatal life stages.

*B. breve* UCC2003 is a model strain that was previously isolated from the stool of a breast-fed infant [28, 29]. Although human-associated, numerous previous studies have shown this strain can efficiently colonise (long-term) the mouse gastrointestinal tract, which we also observed in this study [30, 31]. Importantly, although at lower levels (~10^5^ CFU/g), we observed *B. breve* UCC2003 within the small intestine, linking to our subsequent observations of significant impacts on the IEC transcriptome from this intestinal region. Furthermore, our microbiota profiling suggests minimal impacts on the wider microbiota (at genus level) after supplementation, suggesting that *B. breve* UCC2003 is principally driving specific transcriptomic outcomes. However, we cannot discount that *B. breve* is driving more nuanced microbiota changes, which may also be contributing to downstream IEC responses.

*B. breve* is known to confer beneficial effect on gut health, however our knowledge related to the mechanisms underlying these responses are limited. Most studies have focused on targeted immune cells or pathways (during disease and/or inflammation), and to our knowledge no studies have probed global transcriptomic changes within IECs - the frontline physical barrier between bacteria and host [32, 33]. Our presented findings: ~4,000 up-regulated DEGs and ~450 down-regulated DEGs within the *B. breve* group indicate that this *Bifidobacterium* strain modulates whole-scale changes within this critical single cell layer. Notably, we also examined how *B. breve* modulates adult IEC responses, however, we did not observe any significantly differentially regulated genes when compared to control animals. The striking differences in DEGs between these two life points indicates that *B. breve*-modulation of IECs is limited to the neonatal window. Dominance of *Bifidobacterium* in early life (including strains of *B. breve*) overlaps with the development and maturation of many host responses, including epithelial barrier integrity. Therefore, presence of these strains would be expected to play an over-sized role in this initial homeostatic priming, that may afford protection against inflammatory insults in later-life, as has been shown previously in a mouse model of pathological epithelial cell shedding [17].

Exploring the transcriptional responses in more detail revealed that expressions of key genes associated with formation of epithelial barrier components were up-regulated, including major cell junction protein encoding genes (75%; 42/56 genes). More specifically, several integrin-associated genes were up-regulated in the presence of UCC2003. Integrins facilitates cell-cell and cell-extracellular matrix ECM adhesion and binding, and assembly of the fibronectin matrix that is pivotal for cell migration and cell differentiation [34-36]. Integrins also play an important role in downstream intracellular signalling that controls cell differentiation, proliferation and cell survival, including the Raf-MEK-ERK signalling pathway (we also observed enrichment of genes involved in this pathway) [37, 38]. Another key intestinal barrier component is represented by tight junctions; linking complexes between intercellular spaces, and comprise transmembrane proteins including occludins, claudins, zona occludens and junctional adhesion molecules [13, 39]. Dysfunctional tight junctions may lead to a ‘leaky’ gut, which is characteristic of numerous intestinal disorders including inflammatory bowel diseases [40]. Notably, previous work has suggested early life microbiota disruptions (via antibiotic usage) and reductions in *Bifidobacterium* are correlated with increased risk and/or symptoms of ulcerative colitis and Crohn’s disease [41-45]. A wide range of TJ-related genes were up-regulated after UCC2003 supplementation, particularly *Tjp1* (that encodes ZO-1), *Jam2* and *Claudin34c1*, with a previous study indicating other *Bifidobacterium* species (i.e. *B. bifidum)* also modulate TJ expression via ZO-1 [46]. These data indicated that specific strains of *Bifidobacterium* may modulate key barrier integrity systems during the neonatal period, and therefore absence of this key initial bacterial-host crosstalk may correlate with increased risk of chronic intestinal disorders in later-life [44]. Intestinal mucus, encoded by *Muc* genes (up-regulated due to *B. breve* UCC2003 in this study), plays a crucial role in colonic protection via formation of a physical barrier between the gut lumen and IECs, and deficiencies in MUC-2 has been linked with experimental colitis and increased inflammation in IBD patients [47, 48]. We have also observed that *B. breve* UCC2003 significantly increases goblet cell numbers and mucus production (in gnotobiotic and SPF mice; data not shown). Although the mucus layer may impact direct *Bifidobacterium*-IEC interactions, previous studies have indicated that *B. breve* UCC2003 surface molecules, such as EPS and the Tad pilus may modulate IEC function via signaling through TLRs [17, 49]. Moreover, bifidobacterial metabolites, such as short-chain fatty acids may also act to modulate the IEC transcriptome, with previous studies indicating enhanced expression of TJs and cadherins via acetate [9, 14, 50, 51].

Further network and functional analysis indicated clusters of up-regulated DEGs were associated with cell maturation and cell differentiation (as confirmed by cell type specific analysis), suggesting neonatal *B. breve* exposure positively modulates IEC cell differentiation, growth and maturation. Somewhat surprisingly, we did not observe the same type of striking responses in immune pathways, which may be masked by the sheer number of DEGs involved in cellular differentiation and processes. However, pathways such as Toll-like Receptor TLR1 or TLR2 pathways do appear to be enriched (cluster 2 of signalling network analysis). This may link to previous work indicating that the UCC2003 EPS directly signals via TLR2 to induce MyD88 signalling cascades to protect IECs during intestinal inflammation [17]. *B. breve* M-16V was also shown to interact with TLR2 to up-regulate ubiquitin-editing enzyme A20 expression that correlated with increased tolerance to a TLR4 cascade in porcine IECs, further supporting the involvement of *B. breve* in programming key host immunoregulation receptors [52].

Cell type specific analysis of DEGs revealed stem cells as the IEC type most affected by *B. breve*, with absorptive enterocytes least affected despite being most accessible to bacteria in the gut. This implies that *B. breve* or their secreted metabolites can reach the crypts of the small intestinal epithelium. This has been previously suggested by *in situ* hybridisation histology *in vivo* and by *Bifidobacterium*-conditioned media altering the expression of hundreds of host epithelial genes linked to immune response, cell adhesion, cell cycle and development in IECs *in vitro* [17, 53]. All but two of the 37 differentially expressed stem cell marker genes were up-regulated in the presence of *B. breve* UCC2003, indicating an activating effect resulting in increased pluripotency of stem cells, increased quantity of stem cells and/or an increased quantity of semi-differentiated cells. Single cell sequencing of IECs could be used to further investigate this finding. Thirty-two TFs were predicted to regulate these stem cell signature genes, providing possible targets for future investigation of the mechanisms underlying these responses. Functional analysis of the stem cell signature genes and their regulators suggests *B. breve* increases pluripotency of stem cells and/or semi-differentiated epithelial cells through WNT signalling and nuclear hormone signalling [54]. Furthermore, the overrepresentation of the process “RUNX1 regulates transcription of genes involved in differentiation of HSCs” indicates a possible role for histone methylation in response to *B. breve* UCC2003 [55]. Further determination of host metabolome and proteome after *B. breve* exposure may allow identification of the specific underlying molecular mechanisms [53].

In conclusion, *B. breve* UCC2003 plays a central role in orchestrating global neonatal IEC gene responses in a distinct manner; modulating genes involved in epithelial barrier development, and driving universal transcriptomic alteration that facilitates cell replication, differentiation and growth, particularly within the stem cell compartment. This study enhances our overall understanding of the benefits of specific early life microbiota members in intestinal epithelium development, with potential avenues to explore for subsequent development of novel live biotherapeutic products. Further work exploring time-dependent transcriptional responses, impact of other *Bifidobacterium* species and strains (and use of mutant strains), in tandem with metabolomic and proteomic approaches are required to fully understand the key host pathways and bifidobacterial molecules governing development and maturation of the intestinal barrier during early life.

## Methods

### Animals

All animal experiments and related protocols were performed in accordance with the Animals (Scientific Procedures) Act 1986 (ASPA) under project licence (PPL: 80/2545) and personal licence (PIL: I68D4DCCF), approved by UK Home Office and University of East Anglia (UEA) FMH Research Ethics Committee. C57BL6/J neonatal female mice (n=10) were housed within UEA Disease Modelling Unit and fed autoclaved chow diet ad libitum. Mice were euthanised via ASPA Schedule 1 protocol (CO_2_ and cervical dislocation).

### Bacterial culturing, inoculum preparation and CFU enumeration

*B. breve* UCC2003 (also known as NCIMB 8807) was streaked from frozen glycerol stocks onto autoclaved Reinforced Clostridial Agar (RCA) plates (Oxoid, UK) and incubated in an anaerobic chamber (miniMACS, Don Whitley Scientific) at 37°C for 48 h prior to picking single colonies for inoculation in prewarmed sterilised Reinforced Clostridial Medium (Oxoid, UK).

For preparation of gavage inoculums, 5 ml of inoculated broth was incubated overnight followed by sub-culturing into 5 ml De Man, Rogosa and Sharpe (MRS) medium (Oxoid). After an additional overnight incubation, another sub-culturing into 40 ml RCM was performed. Inoculums were prepared from cultures by 3 rounds of centrifugation at 3220 g for 10 min followed by three PBS washes before dilution in 4 ml (adult mice) or 2 ml (neonatal mice) sterile PBS. Bacterial concentration of inoculum was enumerated by plating serial dilutions in sterile PBS on RCA plates and enumerating colonies following two-day incubation to calculate CFU/ml.

### Bacterial treatment and gut colonisation

Neonatal mice were colonised with *B. breve* UCC2003 by oral gavage with bacterial inoculations of 10^8^ CFU/ml in 50 μl every 24 h for 3 consecutive days. Control mice received oral gavages of sterile PBS. *B. breve* UCC2003 colonisation was confirmed by collection of fresh faeces or intestinal content homogenised with 1 ml sterile PBS followed by serial-dilution plating in sterile PBS on RCA supplemented with 50 mg/L mupirocin and counting of colonies following 2-day incubation to calculate CFU/mg.

### Gut microbiota profiling by 16S rRNA amplicon sequencing and analysis

Genomic DNA extraction of mouse caecal samples on day 4 was performed with FastDNA Spin Kit for Soil following manufacturer’s instructions and extending the bead-beating step to 3 min as described previously [56]. Extracted DNA was quantified, normalised and sequenced on Illumina MiSeq platform using a read length of 2 × 300 bp, sequencing reads were analysed using OTU clustering methods (QIIME v1.9.1) to assign bacterial taxonomy and visualised as described previously [57, 58]. PCA was performed via R package *ggfotify* function *autoplot* and *prcomp*, while diversity index was computed via package *vegan* [59-61]. LDA was performed via LEfSe on Galaxy platform using default parameters [62]. All related graphs were otherwise plotted using R package *ggplot2* [63].

### Tissue collection and isolation of small intestinal epithelial cells (IECs)

Upon tissue harvesting, 0.5 cm sections of small intestines were collected and incubated in 200 μl RNAlater^™^ (Thermo Fisher Scientific) at 4°C for 24 h. Samples were removed from RNAlater^™^ following incubation, blotted dry on filter paper and stored at −80°C until further analysis. An adapted Weisser method was applied for isolation of IECs [17]. Sections (10cm) of small intestines were collected in ice-cold PBS, dissected into 0.5 cm2 pieces and placed in 200 ml Duran bottles. Faecal matter was washed off by inverting 10 times in 0.154M NaCl and 1mM DTT. Liquid was drained and mucus layer removed through incubation of samples in 1.5mM KCl, 96mM NaCl, 27 mM Tri-sodium citrate, 8mM NaH_2_PO4 and 5.6mM Na2HPO4 for 15 min at 220 rpm and 37 °C. IECs were separated from basal membrane by incubation in 1.5 mM EDTA and 0.5 mM DTT for 15 min at 200 rpm and 37 °C followed by shaking vigorously 20 times. IECs were collected from solution by centrifugation at 500 g for 10 min at 4 °C. Supernatant was then discarded and cell pellet resuspended in 3 ml of ice-cold PBS. Cell concentrations of isolated IEC samples calculated by labelling dead cell with trypan blue at a 1:1 v/v ratio and enumeration of viable cells using a Neubauer haemocytometer on an inverted microscope (ID03, Zeiss).

### RNA extraction and sequencing

RNA was extracted from IEC isolations by adding a volume containing 2 x 10^6^ cells in PBS to QIAshredder spin columns (QIAGEN) followed by centrifugation at 9,300 g for 1 min. Follow-through was mixed with 600 μl RLT lysis buffer and used for subsequent RNA isolation. Homogenised sample in RLT buffer from both tissue and IEC isolations were processed by adding 700 μl of 70% ethanol and mixing by pipetting. Sample was then added into RNeasy spin column and spun at 8,000 g for 15 sec. Flow through was discarded and process repeated until all of sample was filtered through column. Then 700 μl of buffer RW1 was added to column and centrifuge at 8,000 g for 30 s. Again, flow through was discarded and filter placed in a new collection tube. To the filter, 500 μl RPE was added and spun at 8,000 g for 30 s followed by discarding of flow through. An additional 500 μl RPE was pipetted into column and centrifuged at 8,000 g for 2 min. Spin column was then placed in a new collection tube and centrifuged at 8,000 g for 2 min. Columns were transferred to a RNA low-bind Eppendorf tube and 30 μl of RNase free water added to directly to the filter. After an incubation of 1 min at RT, sample was centrifuged at 8,000 g for 1 min and flow through containing RNA stored at −80°C.

Purified RNA was quantified, and quality controlled using RNA 6000 Nano kit on a 2100 Bioanalyser (Agilent). Only samples with RIN values above 8 were sequenced. RNA sequencing was performed at the Wellcome Trust Sanger Institute (Hinxton, UK) on paired-end 75 bp inserts on an Illumina HiSeq 2000 platform. Isolated RNA was processed by poly-A selection and/or Ribo-depletion. All samples were sequenced using non-stranded, paired-end protocol.

### Sequence pre-processing and Differential Gene Expression (DGE) analysis

Sequencing quality of raw FASTQ reads were assessed by FastQC software (v0.11.8). FASTQ reads were subsequently quality-filtered using fastp v0.20.0 with options -q 10 (phred quality <10 was discarded) followed by merging reads into single read file for each sample (merge-paired-reads.sh) and rRNA sequence filtering via SortMeRNA v2.1 based on SILVA rRNA database optimised for SortMeRNA software [64, 65]. Filtered reads were then unmerged (unmerge-paired-reads.sh) and ready for DGE analysis.

Transcript mapping and quantification were performed using Kallisto v0.44.0 [66]. Briefly, *Mus musculus* (C57BL/6 mouse) cDNA sequences (GRCm38.release-98_k31) were retrieved from Ensembl database and built into an index database with Kallisto utility index at default parameter that was used for following transcript mapping and abundance quantification via Kallisto utility quant at 100 bootstrap replicates (-b 100) [67].

Differential Gene Expression (DGE) analysis was performed using R library Sleuth (v0.30.0) [68]. Gene transcripts were mapped to individual genes using Ensembl BioMart database with Sleuth function *sleuth_prep* with option *gene mode = TRUE*. Genes with an absolute log2(fold change) >1.0 and q value <0.05 (or, False Discovery Rate; FDR) were considered to be differentially expressed (or, significantly regulated) [69].

### Functional annotation and enrichment analysis

Functional assignment and enrichment analysis was performed using PANTHER Classification System [70]. Briefly, for functional assignment analysis, a list of genes of interest in Ensembl IDs were supplied to the webserver to be mapped to the Mouse Genome Database (MGD) to generate functional classification on those genes of interest [71]. For functional enrichment analysis, a gene list was supplied together with a background gene list in Ensembl IDs to Panther web server, then selected ‘functional overrepresentation test’ and chose a particular function class in the drop-down menu. Recommended by the database developers, Fisher’s exact test and False Discovery Rate (FDR) correction were used to perform enrichment analysis [72]. FDR <0.05 was used as the default cut-off for significant enrichment. Graphs were plotted in R using ggplot2 library [61]. Functional annotation of top 20 up/down-regulated genes was assigned manually via Ensembl and/or MGI (Mouse Genome Informatics) databases [71, 73].

### Network, cluster and signalling pathway analysis

A signalling network of all up-regulated DEGs and their first neighbours was built using all available biological signalling databases in the Cytoscape (v3.7.2) OmniPath app (v1, *Mus musculus*) [74, 75]. Modules of highly connected genes within the signalling network were identified using the MCODE plug-in within Cytoscape [76]. Settings below were applied to obtain clusters in the network: degree cutoff = 3, haircut = true, fluff = false, node score cutoff = 0.5, k-core = 3 and max depth = 100.

The nodes of each individual module were tested for functional enrichment based on both Reactome and PANTHER annotations using PANTHER Classification System as described in previous sub-section [70, 77, 78].

### Enrichment of cell type specific marker genes

Cell type signature gene sets for murine intestinal epithelial cells were obtained from Haber et al. [24]. Both droplet and plate-based results were used. Gene symbols were converted to Ensembl IDs using db2db [79]. Hypergeometric significance calculations were applied to test the presence of cell type specific signatures in the list of differentially expressed genes using all expressed genes as the statistical background (normalised counts > 1 in ≥ 1 sample). Bonferroni multiple correction was applied and any corrected p < 0.05 was deemed significant. Genes with normalised counts > 1 in ≥ 1 sample per condition (*B. breve* UCC2003 treated or control) were used to identify cell type signature genes expressed per condition.

### Key regulator analysis

All mouse transcription factor - target gene interactions with quality scores A-D were obtained from DoRothEA v2 *via* the OmniPath Cytoscape app [26, 74, 75]. A subnetwork was generated consisting of differentially expressed stem cell signature genes and all their upstream TFs which were expressed in the transcriptomics dataset (normalised counts > 1 in ≥ 1 sample). These TFs were further filtered for their relevance in the network. Here all expressed genes and their upstream expressed TFs were extracted from the DoRothEA network. A hypergeometric significance test was carried out on any node with degree ≥ 5 to determine if the proportion of connected nodes which are differentially expressed is higher than in the whole network. Any TF with *P* value < 0.05 following Benjamini-Hochberg correction were deemed significant and used to filter the stem cell signature gene subnetwork. Network visualisation was carried out in Cytoscape [75]. Functional enrichment carried out against Reactome pathways as described in previous sub-sections.

## Ethics Approval

All experiments were performed under the UK Regulation of Animals (Scientific Procedures) Act of 1986. The project licence (PPL 80/2545) under which these studies were carried out was approved by the UK Home Office and the UEA Ethical Review Committee. Mice were sacrificed by CO_2_ and cervical dislocation.

## Data and Code Availability

All raw sequencing reads (both RNA-Seq and 16S rRNA amplicon sequencing) have been uploaded to European Nucleotide Archive (ENA) with accession number PRJEB36661. R scripts for Differential Gene Expression analysis are available on https://github.com/raymondkiu/R_scripts/blob/master/sleuth.R while all other R scripts in data visualization will be available upon request.

## Supplemental Information

Supplemental Information can be found online at

## Acknowledgements

This research was supported in part by the Norwich Bioscience Institutes (NBI) Computing infrastructure for Science (CiS) group through the provision of a High-Performance Computing (HPC) Cluster. L. J. H. is funded by a Wellcome Trust Investigator award (100974/C/13/Z) and together with T. K. by a BBSRC ISP grant for Gut Microbes and Health BB/R012490/1 and its constituent project(s), BBS/E/F/000PR10353 and BBS/E/F/000PR10355. T. K. is also funded by the Genomics for Food security CSP grant from the BBSRC (BB/CSP17270/1). A. T. is supported by the BBSRC Norwich Research Park Biosciences Doctoral Training Partnership (grant BB/M011216/1). D .vS. is supported by Science Foundation Ireland (SFI/12/RC/2273-P1 and SFI/12/RC/2273-P2).

## Author Contributions

Conceptualisation, R. K., L. C. H. and L. J. H.; Methodology, R. K., A. T., L. C. H. and L. J. H.; Software, R. K., A. T. and S. C.; Validation, R. K., A. T., T. K. and L. J. H.; Formal analysis, R. K. and A. T.; Investigation, R. K., A. T., L. C. H. and C. L.; Resources, S. C.; Data curation, R. K.; Writing -original draft preparation, R. K., A. T. and L. J. H.; Writing – review and editing, R. K., A. T., D. vS, T. K. and L. J. H.; Visualization, R. K. and A. T.; Supervision, T. K. and L. J. H.; Project administration, R. K.; Funding acquisition, T. K., D. vS. and L. J. H.

## Figure and Scheme Legends

**Fig. S1.**
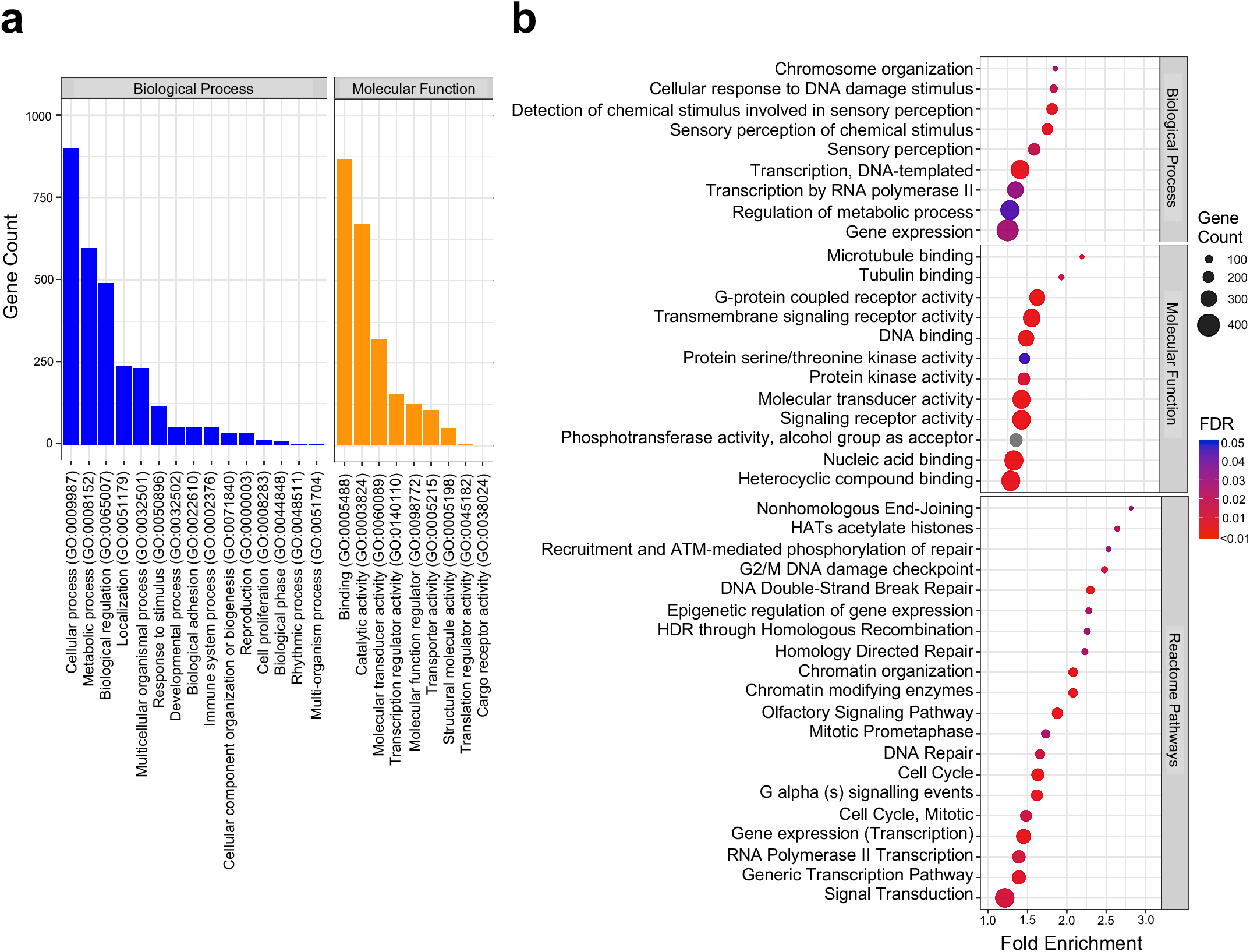
Functional analysis on differentially expressed genes (a) Panther Slim GO-term major categories of significantly up-regulated genes (*n*=3,996). (b) Functional and pathway enrichment analysis on significantly up-regulated genes (Panther Slim GO-term). Only top 20 FDR-ranked enriched pathways (Reactome pathways) are shown. Statistical significance cut-offs: FDR<0.05. Statistical significance: Fisher’s Exact Test. Fold Enrichment was calculated against all expressed genes in IECs as the background (*n*=2l,537).

